# *In vivo* evaluation of *Clostridioides difficile* enoyl-ACP reductase II (FabK) Inhibition by phenylimidazole unveils a promising narrow-spectrum antimicrobial strategy

**DOI:** 10.1101/2023.09.22.559005

**Authors:** Chetna Dureja, Jacob T. Rutherford, Fahad B. A. Pavel, Krissada Norseeda, Isaac Prah, Dianqing Sun, Kirk E. Hevener, Julian G. Hurdle

## Abstract

*Clostridioides difficile* infection (CDI) is a leading cause of hospital-acquired diarrhea, which often stem from disruption of the gut microbiota by broad-spectrum antibiotics. The increasing prevalence of antibiotic-resistant *C. difficile* strains, combined with disappointing clinical trials results for recent antibiotic candidates, underscore the urgent need for novel CDI antibiotics. To this end, we investigated *C. difficile* enoyl ACP reductase (*Cd*FabK), a crucial enzyme in *de novo* fatty acid synthesis, as a drug target for microbiome-sparing antibiotics. To test this concept, we evaluated the efficacy and *in vivo* spectrum of activity of the phenylimidazole analog 296, which is validated to inhibit intracellular *Cd*FabK. Against major CDI-associated ribotypes 296 had an MIC_90_ of 2 µg/ml, which was comparable to vancomycin (1 µg/ml), a standard of care antibiotic. In addition, 296 achieved high colonic concentrations and displayed dosed-dependent efficacy in mice with colitis CDI. Mice that were given 296 retained colonization resistance to *C. difficile* and had microbiomes that resembled the untreated mice. Conversely, both vancomycin and fidaxomicin induced significant changes to mice microbiomes, in a manner consistent with prior reports. *Cd*FabK therefore represents a potential target for microbiome-sparing CDI antibiotics, with phenylimidazoles providing a good chemical starting point for designing such agents.

## INTRODUCTION

*Clostridioides difficile* infection (CDI) is a leading cause of hospital-acquired diarrhea and death (1). The main risk factor for CDI is broad-spectrum antibiotics that disrupt the gut microbiome to facilitate *C. difficile* colonization. Antibiotic treatments for CDI are metronidazole, vancomycin and fidaxomicin, but approximately 20% patients experience recurrent CDI (rCDI) (2–4). During antibiotic therapy, further distortion of the microbiota composition occurs, which increases the risk of recurrence (5–7). Fidaxomicin therapy has a lower recurrence rate than vancomycin and this is thought to result from its narrower spectrum of activity, as it preserves beneficial bacteria that may protect against CDI (7). Resistance to metronidazole, vancomycin and fidaxomicin have emerged, which impacts their therapeutic efficacy (8–10). Furthermore, over the last decade, there have been disappointing clinical trials results for the CDI drug candidates surotomycin and cadazolid (11, 12) and more recently ridinilazole was not found to be superior to vancomycin. Non-antibiotic treatments for CDI have recently been developed, namely bezlotoxumab (13) (a monoclonal antibody to toxin B) and the microbiota transplantation products RBX2660 (14) and SER-109 (15). These non-antibiotics are adjuncts to CDI antibiotics, forming a two-prong approach to treat and prevent rCDI; but microbiota products are not approved for initial CDI episodes. Consequently, there is a need for novel narrow-spectrum CDI antibiotics. However, it is unclear which cellular processes or targets should be pursued to selectively inhibit *C. difficile*, without adversely damaging the microbiome.

Among the known narrow-spectrum drug targets are enzymes that conduct bacterial fatty acid synthesis (16). Bacteria synthesize fatty acids using discrete enzymes in the type II fatty acid synthesis cycle (FASII). The final step of FASII is catalyzed by enoyl-acyl carrier protein (ACP) reductases (ENRs) of which there are four main homologs (FabI, FabK, FabL and FabV) (17). Of these, FabK is a nitronate monooxygenase that uses FMN and NAD(P)H to perform enoyl reductions, whereas FabI, FabL and FabV are short chain dehydrogenases proteins that use only NAD(P)H (17). FabI is a target for antibacterial drugs, as shown by isoniazid that is used for *Mycobacterium tuberculosis* and afabicin that is in phase II clinical trials for *Staphylococcus aureus* infections (18–21). Nonetheless, the physiological relevance of inhibiting FAS-II in gram-positive bacteria was previously debated, on the basis that within an infection bacteria may utilize host lipids to compensate for inhibition of FAS-II (22–24). It is now evident that FAS-II is essential in a species-specific manner. For example, in *S. aureus* the FapR regulator and requirement for branched chain fatty acids prevents this specie from co-opting host lipids to bypass FAS-II inhibition (25, 26). Conversely, in Streptococci and related species the presence of the FabT regulator allows these bacteria to adopt host lipids to circumvent FAS-II inhibition (27, 28). Recently, *C. difficile* FabK (*Cd*FabK) was described as a potential drug target for narrow-spectrum antibiotics for the following reasons (29, 30): the enzyme is conserved in *C. difficile*; FabK is either absent from key gut microbiota or it is carried in combination with FabI; most Clostridia encode FabK along with the FabT regulator, whereas *C. difficile* encodes the FapR regulator (29, 31, 32); and lastly, the FabK enzyme is a druggable target that is amenable to structure-based drug discovery (31–33). Importantly, inhibition of *C. difficile* growth was maintained even in the presence of host lipids, of varying chain lengths and saturation, following genetic silencing of *fabK* or exposure to FAS-II inhibitors (29); this might be due to its carriage of FapR.

Since prior target validation studies for *Cd*FabK were all *in vitro*, which does not entirely replicate the complex milieu in the gut, we conducted *in vivo* studies in mice to further evaluate whether *Cd*FabK is a potential drug target. For this purpose, we used a validated FabK inhibitor 1-[4-(4-bromophenyl)-1H-imidazol-2-yl]methyl-3-[5-(pyridin-2-ylthio)thiazol-2-yl]urea, which is designated herein as 296 (**Fig. 1a**). Compound 296 was first reported to inhibit *S. pneumoniae* FabK (IC_50_ = 0.0017 μM) and lacked inhibition against FabI from *E. coli* (IC_50_>32 μM) or *S. aureus* (IC_50_>100 μM) (30, 34). We reported that 296 also inhibited *Cd*FabK both intracellularly and in enzymatic assays (29). Herein, our *in vivo* studies revealed that 296 was relatively non-absorbed from the gut, it was as efficacious as vancomycin in mice with severe colitis CDI, and it was less disruptive to the gut microbiomes of mice. These findings support *Cd*FabK as a promising drug target and further validates the phenylimidazole class of inhibitors as a promising starting point for designing novel *Cd*FabK inhibitors for therapeutic development.

**Figure 1.**
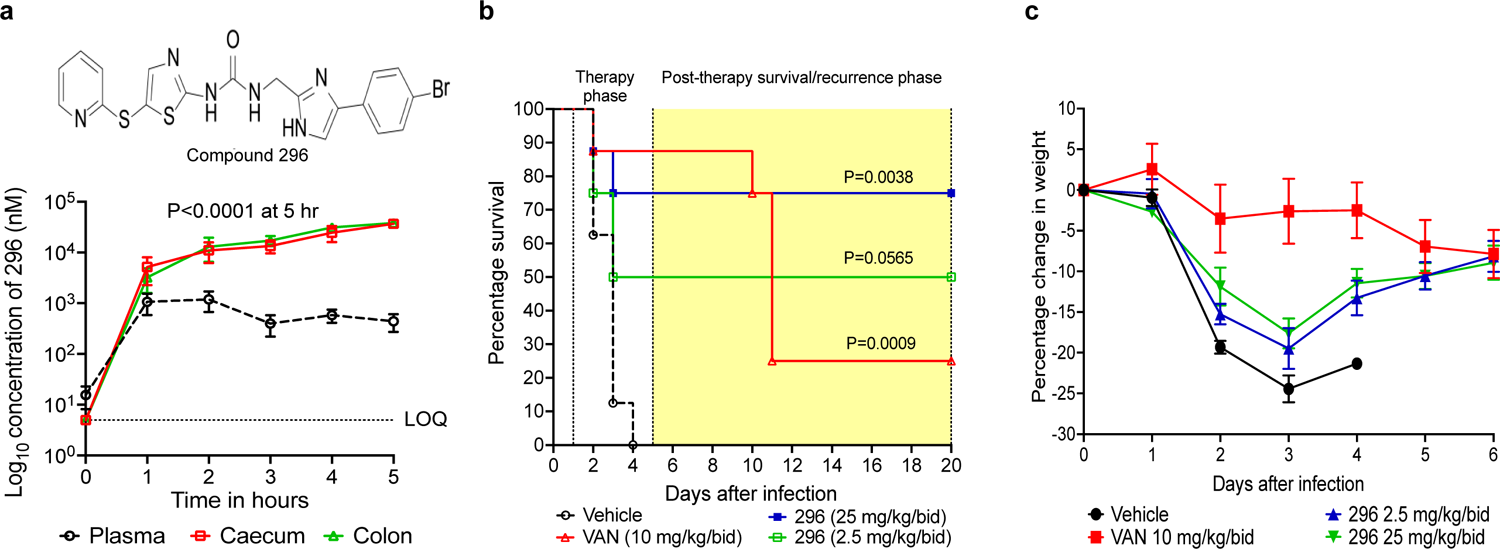
Analysis of 296 *in vivo* pharmacokinetics and efficacy in mice. **(a)** Distribution of 296 in GI tract and plasma; structure of the phenylimidazole 296 is shown. Mice received 100 mg/kg of 296 orally in vehicle (10% DMSO in corn oil). At each time point, 4 mice were culled to measure 296 concentrations by LC-MS/MS. Each data point represents the mean ± SEM from four independent biological samples. **(b-c)** Kaplan-Meier survival analysis and weight changes of mice treated with 296 or vancomycin compared to the vehicle control group. Two experiments were done with total of n=8 mice total (4 males and 4 females). After infection with *C. difficile* R20291 spores, mice were treated 24 hours later, and for 5 days, with vancomycin (10 mg/kg/bid), compound 296 (2.5 mg/kg/bid and 25 mg/kg/bid), or vehicle (10% DMSO in corn oil). Survival is shown in b, whereas changes in weight, as a morbidity symptom, is shown in c for the treatment period (i.e., percentage weight changes are relative to the weight of the mice prior to infection and treatment and are shown as the mean ± SEM). Statistical significances were analyzed in GraphPad prism 10 by one-way ANOVA in a and Log-rank (Mantel-Cox) test in b.

## RESULTS AND DISCUSSION

### CDI-associated ribotypes are susceptible to *Cd*FabK inhibitor

Compound 296 was synthesized as previously described (29, 35) and tested against CDI-associated ribotypes (**Table 1**), including strains that were resistant to metronidazole, vancomycin and fidaxomicin. 296 had MIC_50_ and MIC_90_ values of 0.5 µg/ml and 2 µg/ml, respectively in BHI broth, which were comparable to vancomycin (MIC_50_ and MIC_90_ values of 0.5 µg/ml and 1 µg/ml). It also retained activities (MICs= 1 µg/ml) against a vancomycin resistant strain (MT1470; vancomycin MIC=4 µg/ml) and a fidaxomicin-resistant strain (Modify-28; fidaxomicin MIC= 32 µg/ml). It was found to be bacteriostatic, as determined by MBCs against strain R20291 (epidemic ribotype 027), killing <1 Log of exponential cells (OD_600_nm=0.2) at 64× its MIC of 0.5 μg/ml. In contrast, vancomycin and fidaxomicin were bactericidal; respectively they caused 3-log reductions at 4× and 1× their MICs of 0.5 μg/ml and 0.0625 μg/ml. An eagle effect (36) was observed for vancomycin at 64× its MIC.

**Table 1.** Comparison of activities of 296 and vancomycin against various *C. difficile* isolates.

| *C. difficile* ribotypes<br>(n= number of strains) | MIC (µg/ml) |  |
| --- | --- | --- |
|  | 296 | Vancomycin |
| R20291(027) control | 0.5 | 0.5 |
| 001-072 (n=2) | 0.25-0.5 | 0.5 |
| 002 (n=3) | 1-2 | 0.5-1 |
| 014 (n=3) | 0.5 | 0.25-0.5 |
| 017 (n=2) | 0.125-0.25 | 0.25 |
| 018 (n=1) | 0.5 | 0.5 |
| 019 (n=3) | 1-2 | 0.5-1 |
| 020 (n=3) | 0.062-0.5 | 0.25-0.5 |
| 024 (n=1) | 2 | 0.5 |
| 027 (n=3) | 0.5-1 | 0.25-0.5 |
| 047 (n=1) | 0.125 | 0.25 |
| 054 (n=3) | 0.062-0.5 | 0.25-0.5 |
| 078 (n=2) | 1 | 1 |
| 106 (n=3) | 0.5-1 | 1-2 |
| Range | 0.062-2 | 0.25-2 |
| MIC<sub>50</sub> | 0.5 | 0.5 |
| MIC<sub>90</sub> | 2 | 1 |

### Phenylimidazole 296 is non-absorbed in mice

A desirable property for anti-*C. difficile* antibiotics is that they are non-absorbed, achieving high local concentrations at the site of infection in the large intestine. Previously, *in silico* analysis of phenylimidazole *Cd*FabK inhibitors in QikProp in Schrödinger/Maestro predicted they are low absorption molecules (35). To experimentally test this for 296, we used the Caco2 permeability assay, which also suggested it was non-absorbed with an apparent permeability (Papp) coefficient for apical to basolateral permeability of 21.65 ± 13.57 nm/s, when compared to non-absorbed and absorbed controls vinblastine 5.48 ± 1.66 nm/s and carbamazepine 136.12 ± 50.72 nm/s, respectively. Next *in vivo* pharmacokinetics was examined by giving mice 100 mg/kg of 296 and collecting their intestines and plasma at 1-hour intervals over 5 hours. LC-MS/MS analysis showed that 296 mainly compartmentalized to the ceca and colons of mice (**Fig. 1a**), reaching concentrations of 36,753 ± 1,763 nM and 37,783 ± 1,700 nM in 5 hours, respectively, as compared to plasma (442 ± 170 nM) i.e., ∼83-85-fold higher intestinal concentrations.

### Efficacy of 296 in colitis CDI model

A colitis CDI model (37) was used to evaluate the therapeutic potential of *Cd*FabK inhibition. The model was developed to mimic more hard-to-treat severe CDI development in patients with inflammatory bowel disease (37), which we modified by using dextran sodium sulfate (DSS) at 1.5% w/v instead of 3% w/v and giving antibiotics for 5 days instead of 3 days. Two experiments were done with a total of 8 mice/group (4 males/4 females) that were infected with strain R20291 and 24 hours later they were orally gavaged, twice daily for 5 days with 296 (2.5 or 25 mg/kg), vancomycin (10 mg/kg) or vehicle (10% DMSO in corn oil); vancomycin was administered at a comparable dose used in CDI animal models (38–40). Based on Log Rank statistical analysis of survival over 20 post-infection days, efficacy was observed for all test antibiotics when compared to the vehicle (i.e., the lower and higher doses of 296 had P-values of 0.0565 and 0.0038, whereas vancomycin was P=0.0009) (**Fig. 1b**). Although no statistically significant differences in log rank were observed among the treatment groups, mice that received 296 showed higher overall survival rates following treatment. Specifically, during the post-treatment period (days 5-20) when recurrence occurs, 296-treated mice had survival rates of 50% (2.5 mg/kg/bid) and 75% (25 mg/kg/bid), compared to 25% for vancomycin. Weight changes in mice were also analyzed as a secondary marker of CDI severity and compound efficacy (**Fig. 1c**). Relative weight changes were analyzed, over the 20-day study period, by one-way ANOVA and Tukey’s multiple comparisons post-test in GraphPad prism 10, indicated P values of 0.0453 and 0.0034 for 296 lower and higher doses, respectively and 0.0006 for vancomycin (**Fig. S1** of the Supplementary Materials). While there were no statistically significant differences between the treatment groups, over the full 20 days, mice given 296 experienced more initial weight loss than those given vancomycin and might reflect earlier efficacy by vancomycin. By the fifth day of treatment, 296-treat mice had weights similar to the vancomycin-treated mice.

### Spectrum of activity of 296

Crucial to *Cd*FabK being a drug target is whether its inhibitors are narrow-spectrum. We therefore assembled and MIC tested genome sequenced Gram-positive and Gram-negative gut flora that contained either FabI, FabK, FabI/FabK or homologs of the human FAS-I complex. The panel also included six clostridial species that encode FabK either with FabT (n= 4) or FapR (n= 2). Results showed 296 was inactive against *C. sporogenes* (MICs ≥ 64 µg/ml), but it was active against the other clostridia (MICs=0.5-2 µg/ml) (**Table 2**). Next, to examine if bypass of 296 inhibition occurs in exogenous host fatty acids, we tested *Clostridium histolyticum* (containing a FabT regulator) and *Paeniclostridium sordellii* (containing a FapR regulator) under the same conditions as the controls *C. difficile* R20291 (FapR) and *Streptococcus pyogenes* ATCC 19615 (FabT); cerulenin, an inhibitor of 3-oxoacyl-ACP synthase (FabF) was used as a control. In the absence of 296, all strains grew in the fatty acid mix of stearic acid (0.18 µM), linolenic acid (0.18 µM), myristic acid (0.22 µM) and dodecanoic acid (0.23 µM). 296 retained its activity against the FapR bearing strains i.e., *P. sordellii* (MICs of 1 µg/ml with or without fatty acids) and R20291 (MICs of 1 µg/ml and 2 µg/ml, with or without fatty acids, respectively). Similarly, exogenous fatty acids did not affect the MICs of cerulenin (16 µg/ml versus *P. sordellii*; and 8 µg/ml versus R20291). In contrast, against *C. histolyticum* the presence of fatty acids increased MICs of 296 by 128-fold (i.e., MICs of 64 µg/ml and 0.5 µg/ml, with or without fatty acids, respectively); fatty acids also increased 296 MICs by 16-fold against *S. pyogenes* (MICs of 2 µg/ml and 0.125 µg/ml, with or without fatty acids, respectively). Fatty acids similarly worsened cerulenin MICs against *S. pyogenes* (i.e., 64 µg/ml and 8 µg/ml, with or without fatty acids, respectively; and 8 µg/ml and 1 µg/ml, with or without fatty acids, respectively). These findings accord with reports that FabT may mediate bypass of FAS II inhibition, in contrast to strains whose regulation is mediated by FapR (27, 28). The controls vancomycin and fidaxomicin were inhibitory with MICs of 0.5-4 µg/ml and ≤0.0625 µg/ml, respectively against the clostridial species (**Table 2**), which is consistent with prior reports showing fidaxomicin has excellent activity against most clostridia (41, 42).

**Table 2.**
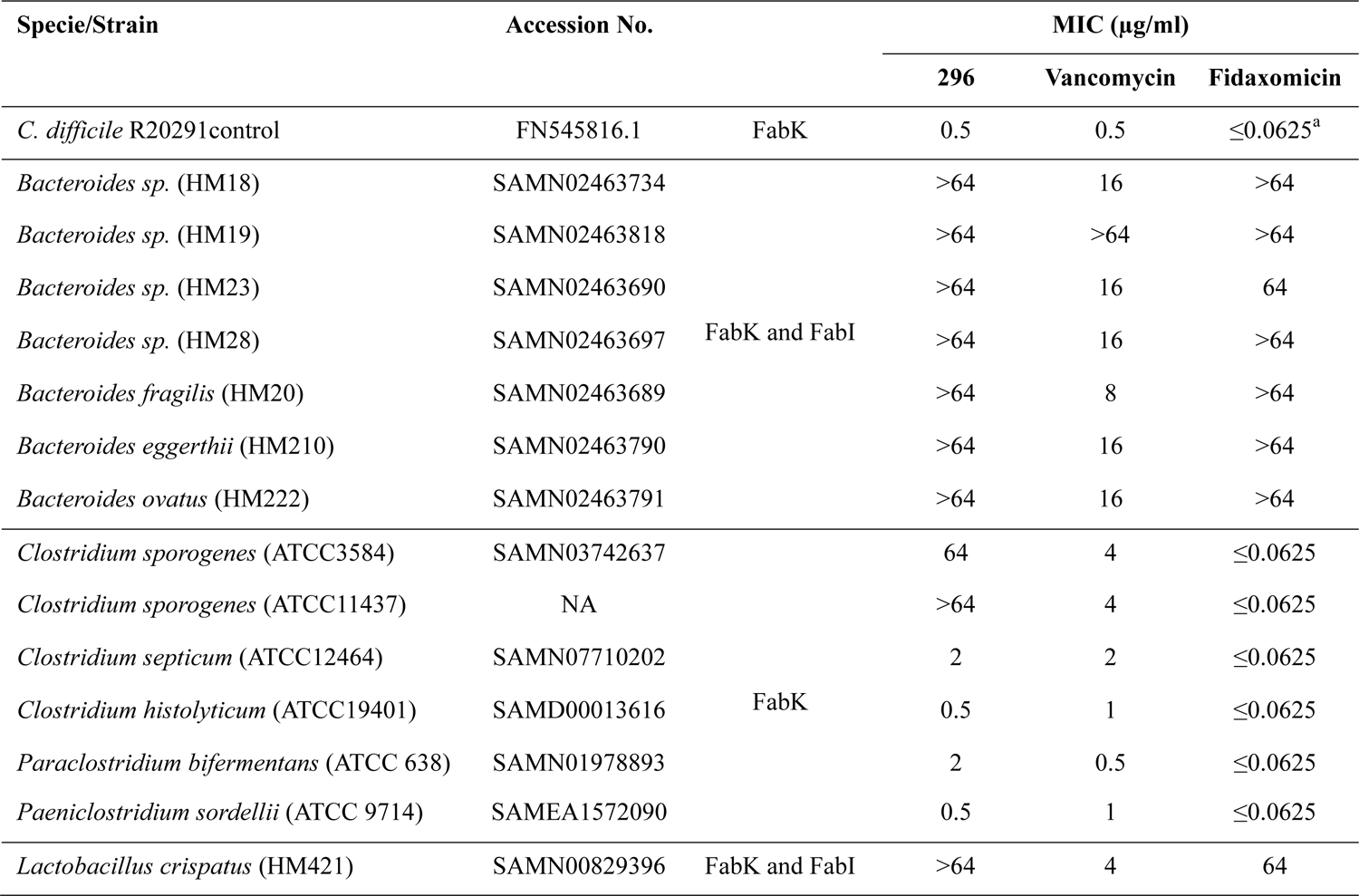
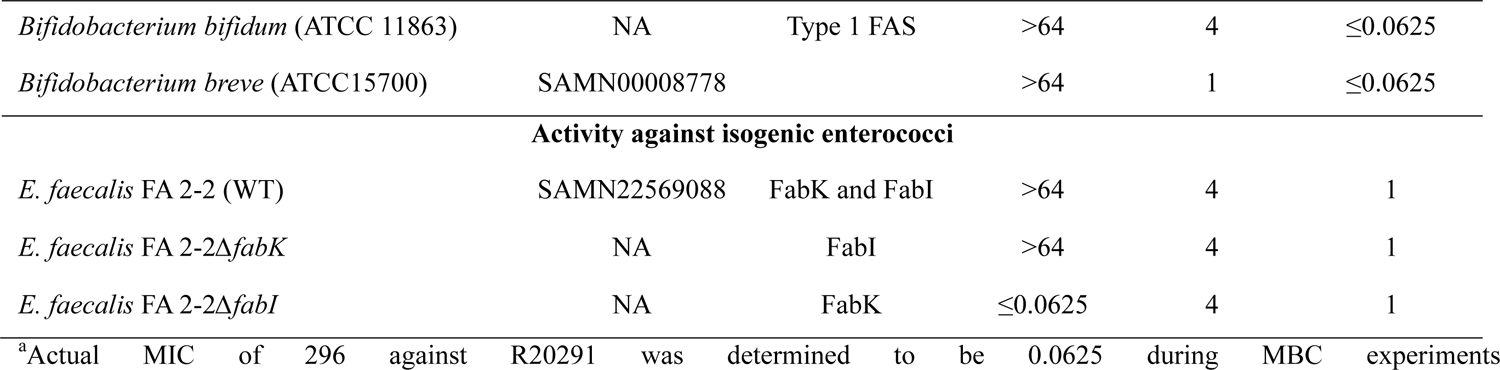
Comparison of MICs of 296 with vancomycin and fidaxomicin against gut species with known enoyl ACP reductases.

Against other genera, 296 was inactive (>64 µg/ml) against *Bacteroides* spp., *Bifidobacterium* spp., and *Lactobacillus crispatus*. Against all non-Clostridia, vancomycin had MICs of 1 to 16 µg/ml, apart from *Bacteroides* sp. HM19 (MIC >64 µg/ml), whereas fidaxomicin was inactive against *Bacteroides* spp. (MIC >64µg/ml) and inhibited *Bifidobacterium* spp. (MIC <0.0625µg/ml). 296 was also inactive against *Enterococcus faecalis*, which is a specie that carries both FabI and FabK. It was reported that *E. faecalis* survives if either FabI or FabK is deleted, although FabI appears to be its preferred ENR (43, 44). Therefore, to test if the activity of 296 is influenced by carriage of ENR, we adopted reported (43) isogenic ENR mutants of *E. faecalis* FA-2, bearing *fabI* or *fabK* (i.e., *E. faecalis* FA-2Δ*fabK* and *E. faecalis* FA-2Δ*fabI*, respectively). 296 was inactive (MICs>64 µg/ml) against wild-type FA-2 and FabI-only mutant but it inhibited growth of the FabK-only strain (MICs≤0.0625 µg/ml; **Table 2**). This further supports that 296 might selectively inhibit organisms bearing FabK, but its activity will depend on the presence of regulators such as FabT that mediates bypass. Against all non-Clostridia, vancomycin had MICs of 1 to 16 µg/ml, apart from *Bacteroides* sp. HM19 (MIC >64 µg/ml), whereas fidaxomicin was inactive against *Bacteroides* spp. (MIC >64µg/ml) but it inhibited *Bifidobacterium* spp. (MIC <0.0625µg/ml).

### 296 did not substantially change gut microbiome diversity

To more holistically measure the effects of 296 on the microbiota species, we adopted a published protocol (**Fig. 2a**) in which mice were given fidaxomicin or vancomycin for three days and stool samples subjected to 16S rRNA analysis (6). We gavaged mice with 296 (25 mg/kg/bid), vancomycin (18.75 mg/kg/bid), fidaxomicin (15 mg/kg/bid) or vehicle (10% DMSO in corn oil) for three days in parallel experiments; the doses of vancomycin and fidaxomicin were previously shown to be physiologically relevant (6, 45). Both drugs were used as controls due to their distinct and differential effects on the microbiota (5, 6).

**(i) Analysis of Shannon diversity.** We first analyzed the 16S rRNA sequences for Shannon diversity (i.e., alpha diversity within samples, in terms of species richness and evenness of the microbiome community). This revealed that 296 did not substantially change the overall microbiome diversity, when compared to the vehicle-treated and pre-treated mice (**Fig. 2b-c**). Moreover, throughout the dosing period, the Shannon diversity for 296 remained comparable to that of the vehicle-treated control (**Fig. 2b**). There was also no statistical difference in the diversities of the vehicle- and 296-treated groups (P=0.908; **Fig. 2c**). In contrast, vancomycin (P<0.0001) and fidaxomicin (P<0.0001) both decreased the overall microbiome diversity, with vancomycin having a more dramatic impact, which is consistent with prior reports (5, 6).
**(ii) Analysis of Beta diversity.** Differences in diversity between samples were evaluated by weighted UniFrac distances and principal coordinate analysis (PCoA). PCoA components 1 and 2 showed a total variance of 50.07% and 27.20%, respectively (**Fig. 3a**). According to the PCoA, over the course of 3 days of treatment, the microbiomes of mice given 296 or the vehicle exhibited close clustering with each other and the baseline pre-treatment group. Conversely, by day 3, the microbiomes of mice given fidaxomicin or vancomycin were significantly dissimilar from each other, and from both the vehicle and pre-treated groups. When considered together, the Shannon diversity and PCoA indicate that 296 minimally disrupted the mouse microbiome.
**(iii) Phyla and class analyses of changes in microbiome composition.** Phyla analysis for 296 and the vehicle groups indicated that after 3 days of treatment, the relative abundance of Firmicutes were 62.9% and 69.8%, respectively, while Bacteroidota were 28.8% and 25.2%, respectively (**Fig. 3b**). These proportions were comparable to pre-treated mice, which had compositions of 67.7% for Firmicutes and 22.4% for Bacteroidota. Conversely, phyla analyses of vancomycin- and fidaxomicin-treated mice showed more significant microbiome composition changes (**Fig. 3b**). Vancomycin enriched Proteobacteria (from 0.4% to 72.5% by day 3) and depleted Firmicutes and Bacteroidota to 10.9% and 0.1%, respectively (**Fig. 3b**). While fidaxomicin also decreased Firmicutes to 12.6%, it significantly increased Bacteroidota to 59.2% (**Fig. 3b**).

**Figure 2.**
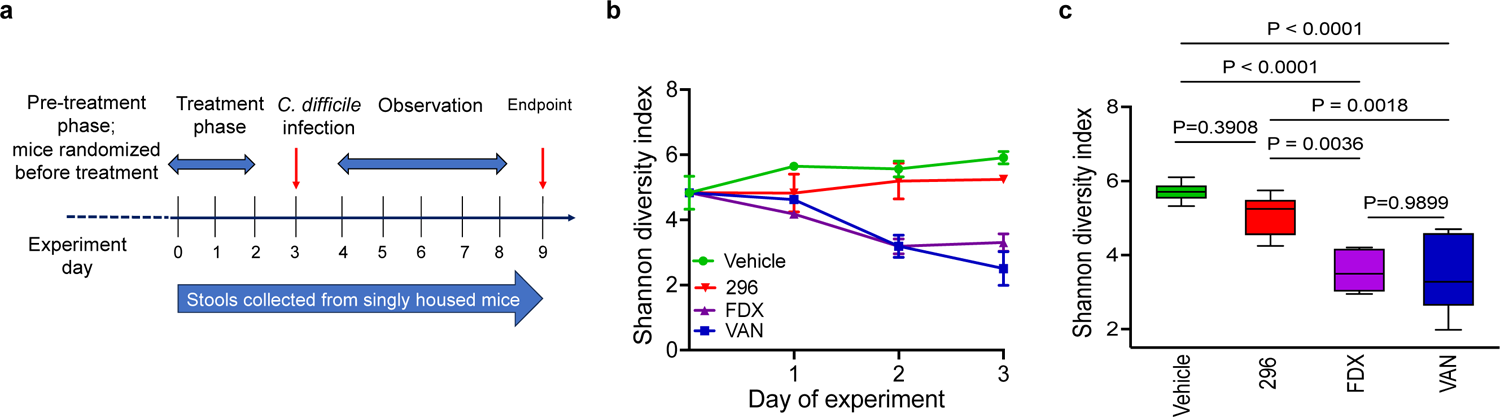
Analysis of effect of 296 on microbiome of mice. **(a)** Schematic of experimental plan of studies in C57BL/6. Prior to treatment, 24 mice (12 males and 12 females) were randomized into 4 groups for treatment with: vehicle (10% DMSO in corn oil), vancomycin (VAN; 18.75 mg/kg/bid), fidaxomicin (FDX; 15 mg/kg/bid), or 296 (25 mg/kg/bid); each randomly assigned mouse was singly housed. On day 0 (pre-treatment), fecal samples were collected, before mice were given the respective treatments for 3 consecutive days. After the treatment phase, mice were infected with *C. difficile* spores. Fecal samples from individually housed male and female mice were pooled for each condition and sequenced for 16S rRNA. **(b)** Operational taxonomic unit-based, within-sample diversity was measured by the Shannon diversity index. Each time point is the mean of two groups (n=6 mice total; 3 male and 3 female mice) and the error bars show the standard error of mean. **(c)** Statistical analysis of the alpha diversity represented in b. The Shannon diversity index of each group on days 1 to 3 (n=6) was combined and statistically evaluated by one-way ANOVA with Tukey’s test in GraphPad prism 10.

**Figure 3.**
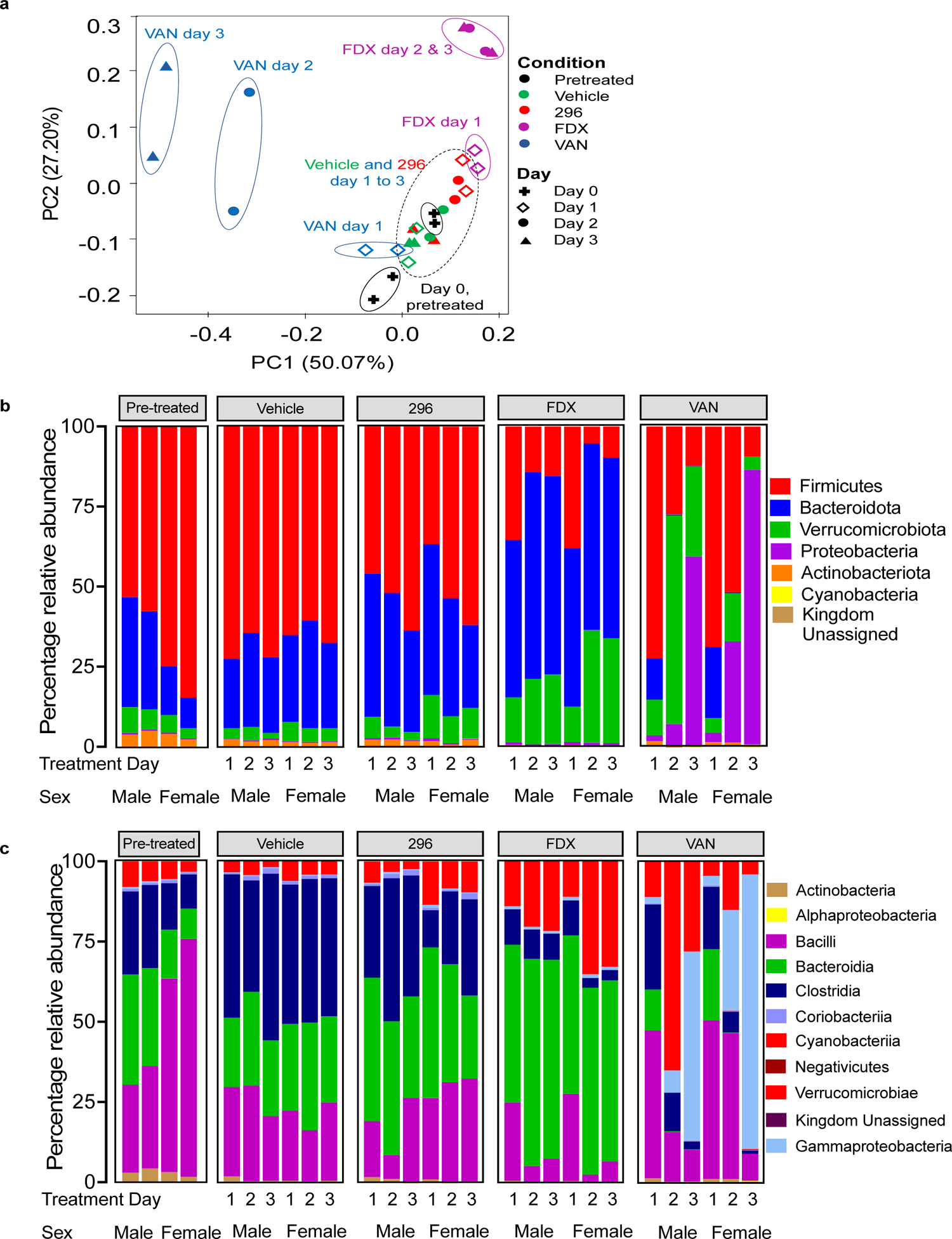
Analysis of changes in microbiome diversity. **(a)** A Principal-coordinate analysis (PCoA), based on weighted UniFrac distances was assessed from 16S rRNA sequence analysis. PCoA 1 and PCoA 2 are shown as a total percentage of the variance of 50.07% and 27.20%, respectively. Each data point in the pre-treated group is from either 6 male or 6 female mice, whereas for the treated sample the data is from pooled fecal samples of 3 mice of either sex. **(b-c)** Stacked bar graph showing the relative abundance of each phylum (b) and class (c) in pre-treated and treated groups. The relative abundance was calculated from the 16S rRNA sequence data.

Across the cohort of pre-treated mice, the three most abundant classes were Bacilli, Bacteroidia, and Clostridia with mean relative abundances of 48.5% ± 22.5%, 22.4% ± 12.0% and 19.2% ± 7.9%, respectively (**Fig. 3c**). This composition is consistent with prior reports in mice (5, 6). It is interesting to note that Bacilli was more dominant in the female mice at pre-treatment, which we are unable to explain. However, Clostridia was the dominant class in the vehicle-treated mice, with relative abundances of average of 44.1%, 39.7% and 47.5% at days 1, 2 and 3, respectively. Bacteroidia and Bacilli had relative abundances of nearly 25% for each class. The proportions of other classes also did not significantly vary during treatment with the vehicle (**Fig. 3c**). In 296-treated mice, the relative abundance of Clostridia was marginally lower than that in the vehicle group over the three days (i.e., 20%, 33.6%, 33.8% on days 1, 2, 3, respectively) (**Fig. 3c**). These findings suggest that 296 may demonstrate partial activity against clostridial species within their natural environment. We speculate that it exhibits selective activity in the gastrointestinal tract. Vancomycin and fidaxomicin caused significant reductions in Clostridia over the treatment period; for vancomycin, the proportions of Clostridia changed from 22.9%, 9.2%, and 1.7% on days 1 to 3, while changes with fidaxomicin were 11.0%, 6.1%, and 5.7%, respectively (**Fig. 3c**). After 3 days of vancomycin, Gammaproteobacteria was the most dominant class, increasing from 2.4% on day 1 to 72.3% on day 3. Conversely, with fidaxomicin, the Bacteroidia class shifted from 49.2% on day 1 to 59.2% on day 3.

### Mice treated with 296 maintained colonization resistance to *C. difficile*

It is widely accepted that a more diverse microbiome enhances resistance to *C. difficile* establishment within the gut community, a phenomenon known as colonization resistance (46). To test this concept, we infected mice with *C. difficile* following three days of compound treatment and monitored fecal burdens for six days post-infection (6). Our results showed that mice treated with the vehicle and 296 had the lowest *C. difficile* bioburdens (**Fig. 4**), compared to mice given fidaxomicin and vancomycin. This suggested that vehicle- and 296-treated mice were more resistant to *C. difficile* colonization, which is consistent with those mice having more diverse microbiomes. Conversely, vancomycin-treated mice displayed the highest susceptibility to *C. difficile* colonization, consistent with having the least diverse microbiome. Our results for vancomycin and fidaxomicin are in line with published findings (6). Based on our findings, it appears that 296 exerts a minimal impact on the microbiome, thus preserving colonization resistance.

**Figure 4.**
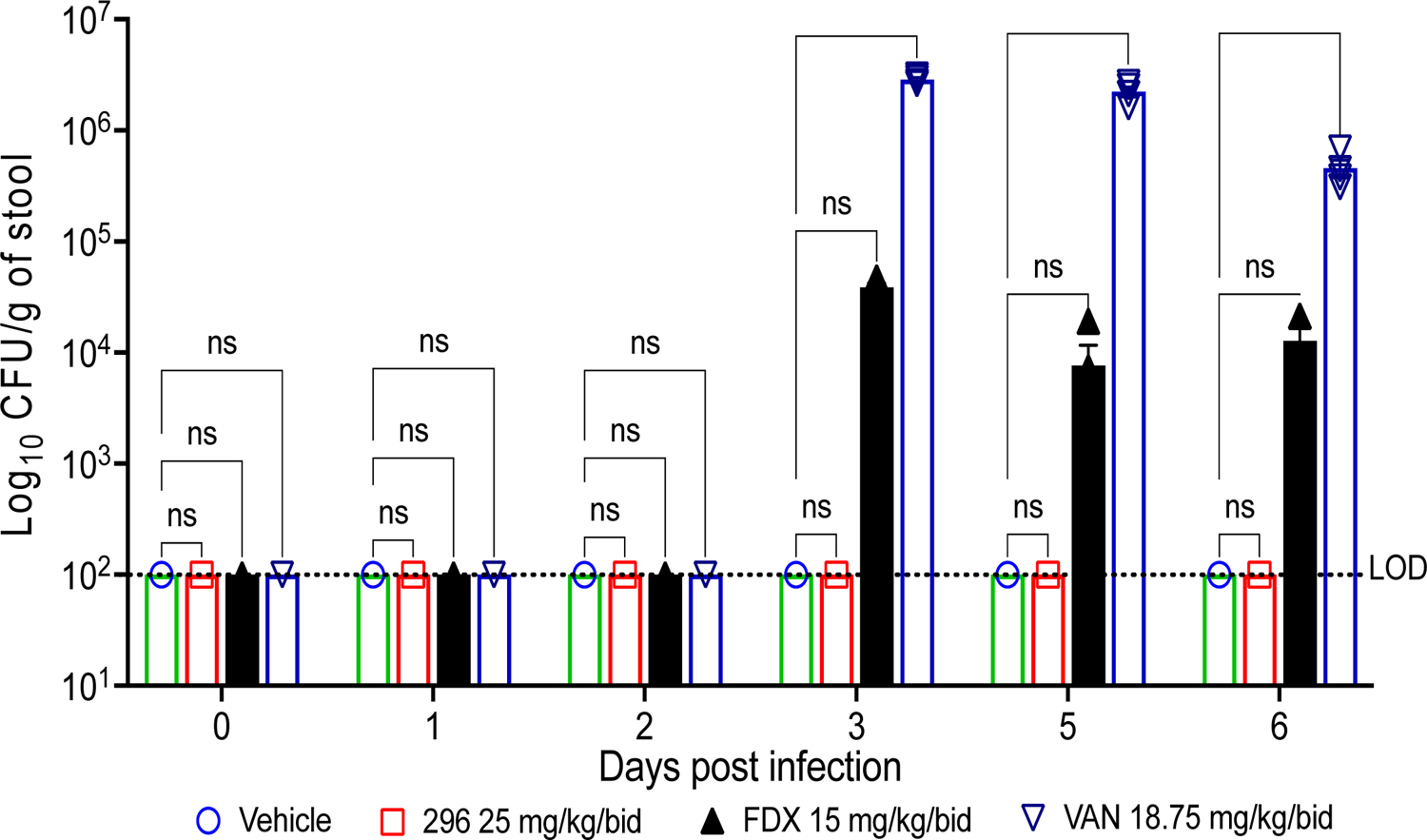
Assessment of colonization resistance following treatment. Mice (3 males and 3 females) were given vehicle (DMSO in 10% corn oil), vancomycin (VAN), fidaxomicin (FDX, and compound 296 for 3 days, before being infected with *C. difficile* R20291 on day 4; each mouse was housed singly. Fecal samples from 3 males or 3 females of each treatment were pooled for each day and *C. difficile* bioburden measured. The data in each bar is from 2 experimental replicates done on different pools and are shown as mean ± SEM; LOD=Limit of detection. Statistical significance was assessed by two-way ANOVA with Dunnett’s test in GraphPad prism 10; ns, not significant; ****, P-value <0.0001.

## CONCLUSION

Narrow-spectrum antibiotics are desirable treatments for CDI, to reduce the likelihood of rCDI onset, as they may promote restoration of normal gut flora that compete with *C. difficile* for resources or they may modify the metabolomic environment to prevent its germination and colonization (46). The benefit of such agents is exemplified by fidaxomicin, whose selective activity results from *C. difficile* RNA polymerase β′ subunit having a specific residue (lysine-84), which forms a salt bridge with the phenolic oxygen of fidaxomicin (47). Lysine-84 is thought to sensitize *C. difficile* to fidaxomicin and its discovery supports the idea that genetic polymorphisms in known drug targets and pathways can be exploited to discover narrow spectrum drugs for CDI. In this study, our *in vivo* microbiome results also suggest that the enzyme *Cd*FabK can be a narrow-spectrum drug target, since the phenylimidazole 296 showed more selective activity in the microbiome of mice, when compared to vancomycin and fidaxomicin. Interestingly, while Clostridial species also bear FabK as their enoyl ACP reductase, mice treated with 296 maintained a high relative abundance of this bacterial class. However, further studies are required to pinpoint species that may become diminished during treatment with a *Cd*FabK inhibitor, in both naïve and disease mice, and the extent to which the microbiota composition is dictated by the type of FAS-II regulation. Although free lipids are known to occur in the intestines of mice, we did not measure free lipids in the intestines of mice in this study.

Phenylimidazole 296 was also efficacious in an acute CDI setting and was associated with an overall higher survival than vancomycin. Although fidaxomicin was not included in these efficacy studies, it is reported to show similar efficacy to vancomycin in mice with acute CDI (48). Hence, we would expect that in mice a *Cd*FabK inhibitor might show efficacy that is comparable to fidaxomicin. We must emphasize that 296 is not a clinical candidate, but it was only deployed as an experimental probe. Nonetheless, from a drug development standpoint, the 296-related class of phenylimidazole *Cd*FabK inhibitors also possess lipophilic characteristics to effectively bind to the hydrophobic catalytic pocket of the enzyme, and this physicochemical property may contribute to poor absorption from the intestinal tract (35). Taken together, the studies conducted herein provide experimental evidence supporting that *Cd*FabK is druggable in an acute CDI setting and that a first-generation inhibitor (296) spares the microbiome and maintains colonization resistance.

## MATERIALS AND METHODS

### Bacterial strains

*C. difficile* R20291 was used as a control strain throughout these studies; other *C. difficile* strains were from BEI Resources, National Institute of Allergy and Infectious Diseases at the National Institutes of Health. Varying anaerobes and *S. pyogenes* used in this study were either from BEI Resources or American Type Culture Collection. *Bifidobacterium bifidum* was maintained in Bifidobacterium broth, whereas *Bacteroides* spp. HM19, HM210 and *Lactobacillus crispatus* (HM421) were maintained in Brucella media supplemented with sheep blood, vitamin K1 and hemin (49). All other strains were maintained in pre-reduced Brain Heart Infusion (BHI) media. Strains were grown at 37 °C in an A35 anaerobic workstation (Don Whitley Scientific).

### Minimum inhibitory concentration (MIC) and minimum bactericidal concentration (MBC) determination

MICs were determined by microbroth dilution in BHI broth using inocula of ∼10^6^ CFU/ml of bacteria and doubling dilutions of compound: 296 (0.25 µg/ml to 64 µg/ml), vancomycin (0.25 µg/ml to 64 µg/ml), and fidaxomicin (0.0625 µg/ml to 64 µg/ml). After 24 hours of anaerobic incubation at 37°C, the MIC was defined as the lowest concentration of compound inhibiting visible growth. MBCs were determined by growing R20291 to OD_600_nm of ∼0.2 and exposing the cultures to 296, vancomycin, and fidaxomicin at concentrations of 0, 0.5, 1, 4, 16, and 64 times their MIC. After 24 hours, serial dilutions were plated onto BHI agars supplemented with 5% w/v charcoal and the MBC defined as lowest concentration killing 3 logs of bacteria relative to time zero.

### Caco-2 permeability assay

Colonic epithelial Caco-2 cells were cultured in Minimum Essential Medium (MEM) containing 10% fecal bovine serum (FBS), 100 units/ml of penicillin, and 100 μg/ml of streptomycin and grown at 37°C in a humidified incubator with an atmosphere of 5% CO_2_. Cells were seeded onto inserts of a 96-well Transwell plate at a density of 0.165×10^5^ cells/insert and cultured in the MEM containing 10% FBS for 7 days. Each monolayer was washed twice with HBSS/HEPES (10 mM, pH 7.4) and the permeability assay initiated by adding each compound (10 μmol/L) to inserts (apical side, A) or receivers (basolateral side, B), and then incubated for 2 h at 37 °C. Compound concentrations in fractions collected from the A and B compartments were assessed by Acquity UPLC/MS with a C18 column (Waters, Milford, MA). Apparent permeability coefficients (Pappa, cm/s) of each compound were calculated from the equation, Pappa=dQ/dt×1/AC*0*, where dQ/dt (μmol/s) is drug flux across the monolayer is, C*0* (μmol/L) is the initial drug concentration on the apical side and A (cm^2^) is the surface area of the monolayer.

### Pharmacokinetics in mice

This experiment was conducted using an animal use protocol approved by Texas A & M University. C57BL/6 mice (6-8 weeks; n=24) from Jackson laboratory were weighed and given 100 mg/kg of 296 (in 10% DMSO in corn oil) by oral gavage. At each time 4 mice (2 male and 2 female) were euthanized by CO_2_ inhalation and blood collected by cardiac puncture. Blood, caeca, and colon were collected after time points of 0, 1, 2, 3, 4, and 5 hours. The plasma was stored at −80°C and the caeca and colon samples were stored at −20°C until quantified by LC-MS/MS as follows. **(i) Standard curves.** Standard curves were generated for each biological sample; for plasma, untreated sample was spiked with 296 at concentrations of 0.002mM to 1mM. As an internal standard, we used a reported analog of 296 i.e., 295 (250 nM); 295 is reported as compound 1a in Jones et al., (30). Compounds were extracted from plasma by vortexing in ice-cold methanol and LC-MS/MS performed on the collected supernatants. For untreated cecum or colon samples standard curves were generated using 50 mg of tissue material. Samples from untreated mice were spiked with 296 at concentrations of 0.001 mM to 1 mM. 295 was added to the samples at 125 nM. Compound extraction was performed in ice-cold methanol in a total volume of 400 µL, by homogenizing samples in a bead mill with 5 to 6 ceramic beads in each vial at a speed of 3.5 m/s for 40 seconds running time with a 20 second interval for total 20 cycles. Supernatants were then analyzed by LC-MS/MS. **(ii) Analysis of samples.** Test samples were similarly prepared as above to extract 296, with 295 added as an internal marker. LC-MS/MS was done using a SCIEX TRIPLE QUAD 5500 tandem mass spectrometer (Framingham, MA) coupled with a SHIMADZU HPLC system (Columbia, MD). An Agilent Eclipse Plus C18 5 µm 2.1×50 mm column was used for compound separation. Mobile phase A, consisting of 95% water and 5% acetonitrile, and mobile phase B, consisting of 100% acetonitrile, were used in a gradient solvent system at a flow rate of 0.5 mL/min. Total run time was 4 minutes. Electrospray Ionization (ESI) using positive mode was used to determine the presence of 295 and 296 and mass spectrometric analysis done using a multiple reaction monitoring mode. The ionization spray capillary voltage was 5500 V, the ion source temperature was 500 °C, and CUR, CAD, GS1, GS2 settings were 20, 8, 50, and 50 PSI, respectively.

### Mouse model of colitis CDI

This experiment was conducted using an animal use protocol approved by Texas A & M University. Mice were administered an antibiotic cocktail of kanamycin (0.4 mg/mL), colistin (850 U/mL), metronidazole (0.215 mg/mL), vancomycin (0.045 mg/mL), gentamicin (0.035 mg/mL), and dextran sulfate sodium (DSS 1.5%) in water for 5 days. They were then switched to regular drinking water for 2 days, before receiving clindamycin (10 mg/kg, via intraperitoneal injection). After 24 hours, mice were inoculated with ∼10^5^ viable spores of R20291 via oral gavage. On the day of infection (day 0), mice were randomly assigned to four groups. A day after infection, mice were treated with vancomycin (10 mg/kg/bid), 296 (2.5 mg/kg/bid or 25mg/kg/bid), and vehicle (10% DMSO in corn oil) for 5 days via oral gavage. After infection, mice were assessed daily for changes in weight, temperature, and appearance. Moribund mice were euthanized by CO_2_ inhalation. Two experiments were done concurrently in which mice were infected with two biologically independent spore preparations of R20291.

### Effects of compounds on microbiome and colonization resistance in mice

This experiment was conducted using an animal use protocol approved by Texas A & M University. C57BL/6 mice (6-8 weeks; 6/group) were randomly assigned to four treatment groups vancomycin (18.75 mg/kg/bid), fidaxomicin (15 mg/kg/bid), 296 (25 mg/kg/bid), and vehicle (10% DMSO in corn oil). Each mouse was housed individually and given compound for 3 days via oral gavage. A day after treatment ended, mice were infected with ∼10^5^ viable spores of *C. difficile* R20291 and were monitored for 6 days post-infection. Fecal samples were collected daily from the beginning of the experiment to assess changes in the microbiome and for measuring *C. difficile* bioburdens. On the last day of the experiment, mice were euthanized, and ceca were collected.

### 16S rRNA gene sequencing for microbiome analysis

Fecal pellets from each mouse were evenly pooled and submitted to Seqcenter (Pittsburgh, US) for 16S rRNA gene sequencing. Genomic DNA was extracted using Zymo Research Quick-16S NGS Library Prep Kit and amplicons made using universal primers (Forward: CCTACGGGDGGCWGCAG, CTAYGGGGYGCWGCAG, and Reverse: GACTACHVGGGTATCTAATCC) that target the 16S V3/V4 regions. Samples were sequenced on V3 MiSeq 622cyc flowcell to generate 2×301bp paired-end reads. The sequences were submitted to NCBI under the bioproject number PRJNA1006031.

### Metagenomic analysis

Raw sequence reads were trimmed of sequence adapters and low-quality reads using bcl-convert. The sequences were then imported into Quantitative Insights into Microbial Ecology version 2 (Qiime2) (50), for taxonomic and diversity analyses. Firstly, primer sequences were removed using cutadapt (51). The demultiplexed paired-end reads were denoised with the dada2 plugin (52). The denoised sequences were then assigned operational taxonomic units (OTUs) using q2-vsearch plugin. The OTUs were identified to the lowest taxonomic level with q2-feature-classifier using Silva 138 99% OTUs full-length sequence database as the reference (53). The OTU table was exported in biom format and subsequently analyzed using Agile Toolkit for Incisive Microbial Analyses (ATIMA), developed at the Alkek Center for Metagenomics and Microbiome Research for relative abundance. Alpha and beta diversity analyses were performed using q2-diversity plugin (54) and the corresponding weighted unifrac PCoA artifact format was exported into qiime2R (https://github.com/jbisanz/qiime2R).

## DATA AVAILABILITY

Data described in this study include MIC, MBC results (shown in Tables 1, 2 and main text); pharmacokinetics of compound 296 and survival of animals infected with CDI are described in Figure 2a and 2b; weight loss is shown for 6 days in Figure 2c (the full data set is shown in Figure S1 of the Supplementary Material); 16S sequence files has been deposited in NCBI under the bioproject number PRJNA1006031.

## ACKNOWLEDGEMENTS

This work was supported by the Office of the Assistant Secretary of Defense for Health Affairs through the Congressionally Directed Medical Research Programs (CDMRP) Peer Reviewed Medical Research Program under Award No. W81XWH-20-1-0296. The opinions, interpretations, conclusions, and recommendations are those of the authors and are not necessarily endorsed by the Department of Defense. J.T.R. acknowledges receipt of funding from 5T32AI055449 Molecular Basis of Infectious Diseases Training Grant from the National Institute of Allergy and Infectious Diseases at The National Institute of Health. We are grateful to Xiang Fu of Analytical Technologies Center, Department of Chemical Biology and Therapeutics and Elizabeth Griffith, Department of Chemical Biology and Therapeutics at St. Jude Children’s Research Hospital for performing the Caco-2 permeability studies. We are grateful for the receipt of strains from John E. Cronan, Department of Microbiology, University of Illinois and BEI Resources, National Institute of Allergy.

